# A Transcriptome Wide Association Study implicates specific pre- and post-synaptic abnormalities in Schizophrenia

**DOI:** 10.1101/384560

**Authors:** Lynsey S Hall, Christopher W Medway, Antonio F Pardinas, Elliott G Rees, Valentina Escott-Price, Andrew Pocklington, Peter A Holmans, James TR Walters, Michael J Owen, Michael C O’Donovan

**Author notes:** joint first author (Lynsey S Hall, Christopher W Medway). Corresponding author: Dr Lynsey Hall MRC Centre for Neuropsychiatric Genetics and Genomics Cardiff University Hadyn Ellis Building Cardiff CF24 4HQ.

## Abstract

Schizophrenia is a complex highly heritable disorder. Genome-wide association studies have identified multiple loci that influence the risk of developing schizophrenia, although the causal variants driving these associations and their impacts on specific genes are largely unknown. Here we link genetic findings to gene expression in the human brain by performing a transcriptome-wide association study (TWAS) in which we integrate the largest published genome-wide association dataset of schizophrenia, with publically available post mortem expression data from the dorsolateral prefrontal cortex (DLPFC). We identify a significant correlation between schizophrenia risk and expression at eighty-nine genes in DLPFC, including forty-two genes not identified in earlier TWAS of this transcriptomic resource. Genes whose expression correlate with schizophrenia were enriched for those involved in nervous system development, abnormal synaptic transmission, reduced long term potentiation, and calcium-dependent cell-cell adhesion. Previous genetic studies have implicated post-synaptic glutamatergic and gabaergic processes in schizophrenia; here we extend this to include molecules that regulate presynaptic transmitter release. We identify specific candidate genes to which we assign predicted directions of effect in terms of expression level, facilitating downstream experimental studies geared towards a better mechanistic understanding of schizophrenia pathogenesis.

## Introduction

Schizophrenia is a severe psychiatric disorder which typically manifests in late adolescence or early adulthood, although prodromal functional impairment may precede formal diagnosis by many years^1^. Treatments for schizophrenia are symptomatic rather than curative; they are effective in many cases, but a large proportion of people with the disorder have a chronic or relapsing and remitting presentation. As a result, schizophrenia is associated with extensive health and social costs, reduced life expectancy (10-20 yrs) and the suicide rate is 12-times higher than in the general population^2–4^.

Schizophrenia is multifactorial but highly heritable. The genetic contribution is polygenic, involving large numbers of risk alleles spanning the full spectrum of possible allele frequencies^5–7^. Genome-wide association studies (GWAS) have identified large numbers of schizophrenia associated risk loci defined by common alleles of small effect size, the most recent study (40,675 cases and 64,643 controls; referred to hereafter as CLOZUK+PGC2) reporting 174 independent association signals representing 145 physically distinct associated loci^5^.

In principle, each genetic locus identified by GWAS provides an opportunity to expose a biological mechanism bridging statistical association and disease. However, in practice, because the sentinel or index associated polymorphism is often in extensive linkage disequilibrium (LD) with large numbers of flanking polymorphisms, a single associated locus often encompasses multiple genes and regulatory motifs. Allied with the possibility that some of the latter may regulate genes that lie beyond the boundaries of the associated region, the imprecision inherent in GWAS makes extracting biological information from these studies challenging. Seeking to address this, a number of approaches have been developed to prioritise causal variants and/or genes at disease loci based on functional annotations of the genome^8^. Broadly speaking, the aims of integrating genomic and functional genomic data are to link genetic associations to specific candidate genes that may be responsible for driving the genetic associations, and to exploit the combined datasets to identify candidate loci that are not detected by GWAS alone.

For many complex disorders, including schizophrenia, for which the common variant associations appear to be largely driven by variation affecting gene expression or splicing rather than nonsynonymous variation^9,10^, there has been considerable interest in integrating GWAS data with genome wide annotations of regulatory polymorphisms defined in appropriate tissues. Several methods, collectively described as transcriptome-wide association studies (TWAS) have been developed to achieve this, including Summary Mendelian Randomisation (SMR)^11^, PrediXcan^12^ and FUSION^13^, some of which have been applied to studies of schizophrenia. Here the principle is to derive genomic predictors of gene expression from samples (not necessarily cases) for which both genomic and gene expression data are available, and to use those predictors to impute gene expression in independent samples (e.g. case-control samples) for which genome-wide association data, but not gene expression data, are available. In contrast to direct case-control studies of gene expression in human brain, by exploiting genetic predictors of expression via transcriptome imputation into cases and controls using genetic data, it is possible to take advantage of the much larger sample sizes that currently comprise GWAS datasets to identify genes whose expression is predicted to differ between cases and controls. As noted previously, an additional advantage over case-control post-mortem brain expression studies is that by inferring expression from genomic information, the TWAS approach is fairly robust to reverse causality^13^.

Applying the FUSION approach to GWAS summary statistics from the second GWAS of schizophrenia published by the Psychiatric Genomics Consortium (or PGC2), Gusev and colleagues^14^ recently identified 44 genes for which there was significant evidence for association between schizophrenia and the genetically imputed *cis*-component of gene mRNA expression in adult *post-mortem* human brain tissue from the dorsolateral prefrontal cortex (DLPFC). The SMR approach has also been used to identify candidate expression changes that may explain genetic findings^11^; and using this method, we reported association between the expression of 22 genes in DLPFC and schizophrenia liability^5^. Large numbers of genes whose mRNA expression in non-brain tissues (e.g. fat, peripheral blood, testis) have also been correlated with schizophrenia^11,14–16^; while there is clear evidence for cross tissue overlap in the effects of gene variants on gene expression, unless supported by findings in brain tissue, caution in interpreting those findings in the context of schizophrenia is required.

Here we report a FUSION analysis based on the largest published schizophrenia GWAS dataset^5^ and publicly available expression data from the Common Mind Consortium study of DLPFC^17^. We restricted our analysis to this single tissue source as enrichments for common variant genomic and transcriptome-wide polygenic association signals are effectively restricted to the brain^5,14^, and therefore associations between schizophrenia and heritable gene expression derived from this tissue have the highest plausibility of being relevant to the disorder. The use of a single source tissue also facilitates downstream gene-set analyses as the heritability of expression for each gene has been derived from the same sized and equivalently powered sample, and therefore expression of each gene has the same opportunity to be correlated with schizophrenia, subject of course to it being expressed in that tissue. This in turn minimizes potential ascertainment bias when including genes in gene-sets. In the current study we report 89 TWAS significant associations between gene expression and schizophrenia, 44 of which have not been previously reported in other schizophrenia TWAS studies which focus on brain tissue(s)^5,14–16^. TWAS associated genes were enriched in processes involved in synaptic development and plasticity, and impaired long term potentiation. Within these significant gene-sets, we identify individual significant candidate genes to which we assign direction of expression changes in schizophrenia. The findings provide strong candidates for experimentally probing the molecular basis of synaptic pathology in schizophrenia.

## Materials and Methods

TWAS was performed using the FUSION software^13^. Briefly, this method requires GWAS summary statistics with directional information (i.e. Z-score and associated allele), reference haplotypes to estimate linkage disequilibrium, and expression weights derived from subject-level expression data in relevant tissue(s). Reference haplotypes and compiled expression weights were obtained from the FUSION website (http://gusevlab.org/projects/fusion/). The analyses presented here use reference haplotypes from the 1000 Genomes European subpopulation and pre-calculated expression weights for the dorsolateral prefrontal cortex (DLPFC). These expression weights are calculated from the *cis*-genetic (+/-500 kilobases from transcription start/stop site) component of gene expression in individuals from the Common Mind Consortium^17^ (n=452). *Cis*-expression is modelled using four regression frameworks -bslmm, blup, enet, and lasso - and polymorphisms contributing to the model explaining the largest proportion of variance explained (*r^2^*) in gene expression were used to generate TWAS test statistics for each gene. Only genes with significantly non-zero (p ≤ 0.05) *cis*-heritable expression were modelled (n=5 420) and thus available for analysis with FUSION.

GWAS summary statistics were acquired from the CLOZUK+PGC2 schizophrenia GWAS meta-analysis^5^ (40 675 cases, 64 643 controls). Summary statistics were filtered prior to use in FUSION using the munge_sumstats.py (v2.7.13) script, distributed as part of the LD score regression package (https://github.com/bulik/ldsc). Given its localized pattern of long range and complex LD, genetic variants falling within the extended MHC region (chr6:28 477 797 - 33 448 354) were removed prior to analysis, leaving 6 414 705 polymorphisms.

TWAS test statistics were calculated over 22 autosomes using the *FUSION.assoc.test.R* script (R 3.4.0) with default parameters. A multiple-testing correction was applied to TWAS results using the Bonferroni method. Where multiple proximal (+/-500kbp) genes reported a significant TWAS association, statistically independent signals were discriminated using the conditional analysis, conditioning on each TWAS significant gene within discrete loci using a 100 000 kilobase window, implemented in the *FUSION.post_process.R* script. Genes which were TWAS significant (p ≤ 9.43×10^-6^) and remained significant after conditional analysis and locus-wide Bonferroni correction (conditional p ≤ 0.05/2N genes tested) were considered statistically independent.

To assess the relationship between TWAS and GWAS findings, TWAS significant loci were mapped to the genomic coordinates for 145 schizophrenia-associated loci from the *CLOZUK+PGC2* schizophrenia meta-analysis^5^, where GWAS loci were defined for each independent tag-SNP using an LD r^2^ > 0.6, and aggregating proximal (+/-250kbp) loci. At each locus containing a TWAS signal which could be attributed to a single gene, locus-wide GWAS statistics were conditioned on imputed gene expression using the *FUSION.post_process.R* script. The change in p-value of the most significant polymorphism within the locus, before and after conditional analysis, was used to evaluate the effect of imputed expression on phenotypic (schizophrenia) association.

To test the temporal differences in gene expression across developmental stages, a Pearson’s rank correlation was calculated for TWAS Z-scores and effect sizes for the prefrontal cortex across 19 developmental periods, ranging from fetal to adult, using data collected by the BRAINSPAN study which had been subjected to the quality control pipeline as described by Gusev *et al*^14^. P-values were corrected for multiple testing using FDR.

A competitive gene-set enrichment analysis was performed using linear regression in R^18^, with absolute TWAS Z-score as the dependent variable and gene-set membership as a linear predictor. This analysis was carried out for 10 significant candidate gene-sets and 6 677 data-driven gene-sets reported by Pardiñas and colleagues. Further details about these gene-sets can be found here^5,19^. Gene-sets containing fewer than ten cis-heritable genes were removed (resulting in 2 316 data-driven gene-sets). Gene-sets were corrected for multiple testing using a 5% false-discovery rate implemented in R^18^. To test whether gene-sets were significantly depleted for genes with *cis*-heritable expression (i.e. TWAS genes), a non-*cis*-heritable gene list was generated by excluding TWAS genes from a background panel of known brain-expressed genes^20^. Differences between the proportion of genes within a given set which were *cis*-heritable versus non *cis*-heritable were assessed using a Χ^2^ test. As this method does not account for the correlation between genes in the way that gene-set analysis using software for SNP-based data (such as MAGMA) does, a quantile-quantile plot of observed versus expected p-values was generated for this analysis (**Supplementary Figure 1**), which demonstrated no inflation of test statistics above the null (λ = 0.94).

## Results

### TWAS Genes

Of 5 382 genes with significant *cis*-heritable expression in the dorsolateral prefrontal cortex (DLPFC), TWAS test statistics were successfully modelled for 5 301 genes using the schizophrenia association summary statistics. After adjusting for multiple testing using the Bonferroni method, a significant TWAS association signal (p ≤ 9.43×10^-6^) was observed for 89 genes. TWAS test statistics for all genes are shown in **Figure 1** and **Supplementary Table 1**. Schizophrenia liability was not significantly biased towards increased or decreased expression, with increased expression being positively correlated with disease risk at just over half of genes (n = 46, sign test p = 0.83). Proximal TWAS significant genes (those within a 500 kilobase window) were aggregated, resulting in 62 discrete regions of the genome (**Supplementary Table 2**). Conditional analyses within each region identified 68 genes with statistically independent TWAS signals (locus-wide Bonferroni p ≤ 0.05). The TWAS signal could be attributed to a single gene at 57 of these loci (**Supplementary Tables 2 and 3**).

**Figure 1.**
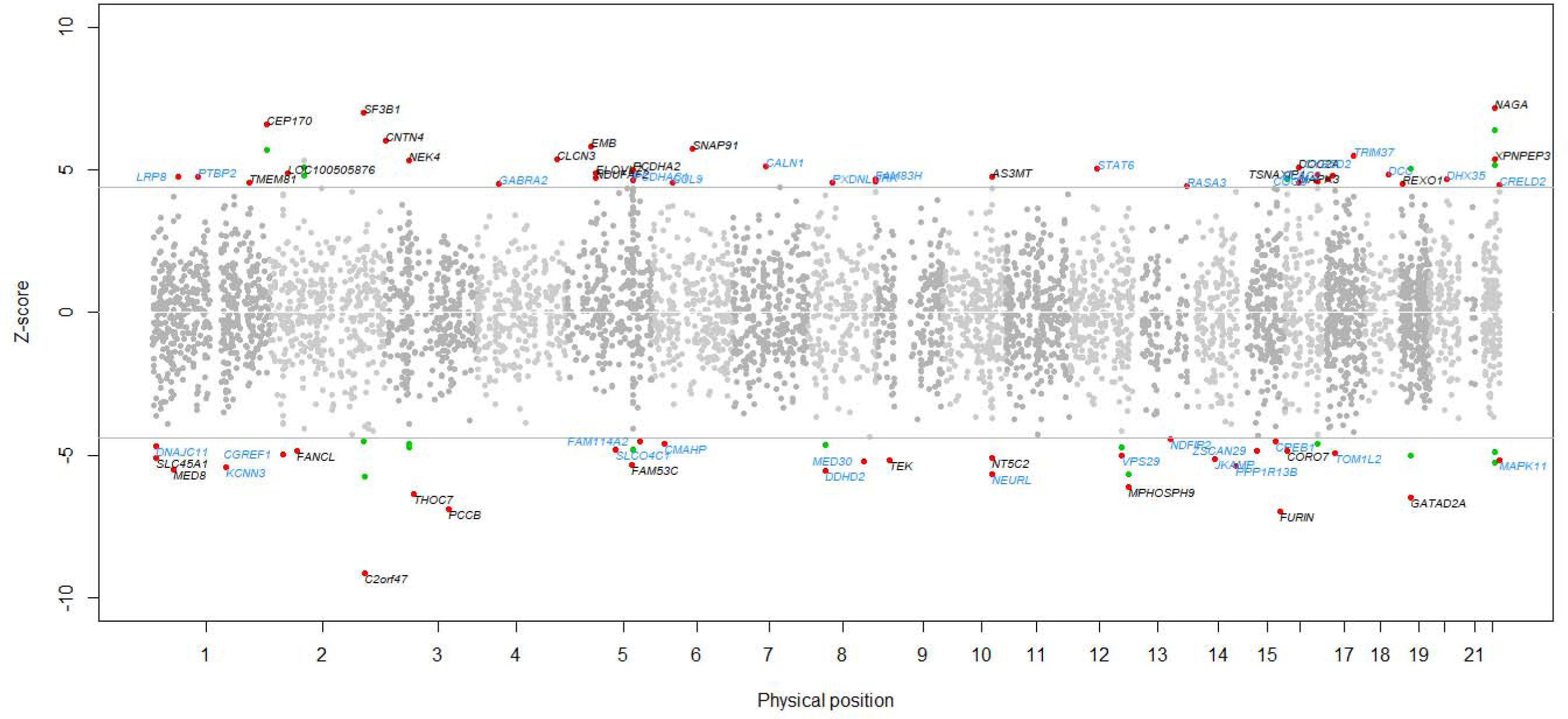
Mirrored Manhattan plot of transcriptome-wide association results. Each point represents a gene, with physical genomic position (chromosome, base-pair) plotted on the x-axis and association Z-score between gene expression in the dorsolateral prefrontal cortex and schizophrenia plotted on the y-axis. Transcriptome-wide-significant associations are highlighted as green points. Conditionally independent significant associations are highlighted as red points, and labelled with gene names in black (if the gene has previously been reported) or blue (if the gene is a novel association).

Thirty-six of the TWAS significant loci, incorporating 56 TWAS significant genes, could be mapped to genome-wide significant loci reported in schizophrenia GWAS upon which this TWAS was based^5^. It was possible to distinguish a single significant TWAS gene at 32 of these loci, after conditioning on other TWAS signals at the locus (**Table 1, Supplementary Table 4**). At 28 of these, conditioning on the imputed expression of the candidate gene nominated by TWAS was sufficient to reduce sentinel SNP associations to below genome-wide significance (p ≥ 5×10^-08^), without unmasking a secondary signal, suggesting that these candidates potentially fully account for the association signals at the locus. Conditional regional association plots for the 32 loci where a single gene was driving the signal are shown in **Supplementary Figures 2 to 33**.

**Table 1:** Thirty-two TWAS significant loci which map to putative schizophrenia risk loci (identified by GWAS), at which the association signal can be attributed to the cis-heritable expression of a single gene in the dorsolateral prefrontal cortex (DLPFC). Locus start and stop denote the locus boundaries as defined in Pardiñas et al schizophrenia GWAS; TWAS test statistics for the association between gene expression in the DLPFC and schizophrenia case/control status are denoted by TWAS Z-score and TWAS P-value. Top GWAS SNP denotes the most significant SNP in the schizophrenia GWAS, followed by its genomic position, GWAS P-value and P-value after conditioning on imputed gene expression. Top cond. SNP denotes the most significantly associated SNP after conditioning on imputed gene expression, its genomic position, GWAS P-value and conditional P-value. Novel gene associations are highlighted in bold.

| Gene | Chr | Locus Start | Locus End | TWAS Z-score | TWAS P-value | Top GWAS SNP | Position | GWAS P-value | Cond. P-value | Top cond. SNP | Position | GWAS P-value | Cond. P-value |
| --- | --- | --- | --- | --- | --- | --- | --- | --- | --- | --- | --- | --- | --- |
| *SLC45A1* | 1 | 8392592 | 8701288 | -5.10 | 3.44E-07 | rs301798 | 8488565 | 5.40E-09 | 4.80E-04 | rs2071414 | 8920712 | 0.003 | 2.80E-04 |
| *MED8* | 1 | 44029353 | 44137257 | -5.50 | 3.73E-08 | rs11210892 | 44100084 | 1.60E-11 | 2.40E-05 | rs11210892 | 44100084 | 1.60E-11 | 2.40E-05 |
| *CEP170* | 1 | 243503764 | 243685760 | 6.59 | 4.51E-11 | rs14403 | 243663893 | 1.70E-10 | 4.60E-06 | rs14403 | 243663893 | 1.70E-10 | 4.60E-06 |
| *FANCL* | 2 | 58386377 | 58468515 | -4.86 | 1.20E-06 | rs11682175 | 57987593 | 3.10E-12 | 4.80E-07 | rs848293 | 58382490 | 8.40E-12 | 3.10E-08 |
| *ALMS1P* | 2 | 73872045 | 73912694 | 5.35 | 9.01E-08 | rs11899902 | 73631564 | 3.10E-09 | 1 | rs2077586 | 73161551 | 3.00E-08 | 4.70E-08 |
| *SF3B1* | 2 | 198256697 | 198299771 | 7.01 | 2.40E-12 | rs699319 | 198281776 | 1.20E-12 | 1 | rs9677260 | 197327735 | 8.60E-06 | 1.40E-07 |
| *C2orf47* | 2 | 200820039 | 200828847 | -9.12 | 7.44E-20 | rs769949 | 200729558 | 4.50E-16 | 0.54 | rs1451488 | 199990107 | 4.70E-12 | 6.00E-10 |
| *CNTN4* | 3 | 2140549 | 3099645 | 6.03 | 1.65E-09 | rs17194490 | 2547786 | 3.20E-12 | 1.50E-05 | rs17014863 | 2519703 | 3.20E-12 | 7.20E-06 |
| *NEK4* | 3 | 52744795 | 52804965 | 5.34 | 9.25E-08 | rs1080500 | 53175017 | 2.70E-12 | 1.80E-06 | rs1080500 | 53175017 | 2.70E-12 | 1.80E-06 |
| *THOC7* | 3 | 63819545 | 63849597 | -6.37 | 1.91E-10 | rs704367 | 63868777 | 2.60E-10 | 0.09 | rs935786 | 63342866 | 4.00E-05 | 5.40E-06 |
| *PCCB* | 3 | 135969166 | 136056737 | -6.89 | 5.39E-12 | rs7432375 | 136288405 | 4.10E-12 | 1 | rs1381251 | 136947138 | 0.08 | 7.00E-05 |
| *CLCN3* | 4 | 170541671 | 170644338 | 5.36 | 8.36E-08 | rs10520163 | 170626552 | 2.80E-08 | 1 | rs13145956 | 170854932 | 0.001 | 9.00E-05 |
| *EMB* | 5 | 49692030 | 49737234 | 5.82 | 5.84E-09 | rs1423645 | 49836715 | 3.00E-08 | 0.66 | rs1363846 | 50280890 | 0.005 | 0.02 |
| *FAM53C* | 5 | 137673223 | 137685418 | -5.32 | 1.04E-07 | rs3849046 | 137851192 | 7.10E-10 | 1.70E-05 | rs3849046 | 137851192 | 7.10E-10 | 1.70E-05 |
| *CMAHP* | 6 | 25093741 | 25166793 | -4.61 | 4.10E-06 | rs7749823 | 26158079 | 6.10E-30 | 1.20E-26 | rs7749823 | 26158079 | 6.10E-30 | 1.20E-26 |
| *SNAP91* | 6 | 84262604 | 84419127 | 5.76 | 8.45E-09 | rs217287 | 84407466 | 9.50E-13 | 1.60E-05 | rs628629 | 83894382 | 2.70E-08 | 3.50E-06 |
| *DDHD2* | 8 | 38089008 | 38120287 | -5.53 | 3.23E-08 | rs7845911 | 38135412 | 7.00E-10 | 0.006 | rs6999014 | 38718619 | 9.60E-05 | 1.90E-05 |
| *JRK* | 8 | 143738873 | 143751401 | 4.59 | 4.40E-06 | rs4129585 | 143312933 | 9.20E-18 | 5.70E-18 | rs4129585 | 143312933 | 9.20E-18 | 5.70E-18 |
| *STAT6* | 12 | 57489186 | 57522883 | 5.05 | 4.34E-07 | rs324015 | 57490100 | 1.40E-10 | 2.10E-05 | rs12814239 | 57569478 | 3.00E-07 | 5.20E-06 |
| *VPS29* | 12 | 110929329 | 110939916 | -5.02 | 5.13E-07 | rs4766428 | 110723245 | 2.70E-14 | 4.60E-09 | 110723245 | 110723245 | 2.70E-14 | 4.60E-09 |
| *MPHOSPH9* | 12 | 123640945 | 123717658 | -6.13 | 9.06E-10 | rs4460848 | 123677810 | 8.80E-16 | 1.50E-07 | rs4460848 | 123677810 | 8.80E-16 | 1.50E-07 |
| *NDFIP2* | 13 | 80055258 | 80130212 | -4.45 | 8.62E-06 | rs3742119 | 79916994 | 2.10E-08 | 5.60E-04 | rs4391942 | 80605356 | 8.20E-04 | 1.10E-04 |
| *JKAMP* | 14 | 59951160 | 59972081 | -5.13 | 2.94E-07 | rs1253107 | 59952947 | 1.50E-07 | 0.12 | rs17097273 | 60743546 | 0.002 | 2.60E-04 |
| *PPP1R13B* | 14 | 104200087 | 104313927 | -5.40 | 6.66E-08 | rs8017993 | 104047734 | 1.50E-13 | 1.30E-07 | rs8017993 | 104047734 | 1.50E-13 | 1.30E-07 |
| *CPEB1* | 15 | 83211950 | 83316728 | -4.53 | 5.81E-06 | rs783540 | 83254708 | 8.50E-10 | 3.50E-05 | rs7163848 | 83398315 | 2.40E-05 | 2.40E-06 |
| *FURIN* | 15 | 91411884 | 91426687 | -6.96 | 3.39E-12 | rs17514846 | 91416550 | 2.70E-12 | 0.02 | rs12899811 | 91544076 | 0.001 | 4.30E-04 |
| *TSNAXIP1* | 16 | 67840976 | 67861971 | 4.86 | 1.20E-06 | rs1975802 | 68285847 | 3.60E-08 | 0.009 | rs2305811 | 68321859 | 3.90E-05 | 0.002 |
| *TOM1L2* | 17 | 17746821 | 17875784 | -4.93 | 8.09E-07 | rs4925114 | 17711270 | 2.60E-08 | 0.01 | rs7223686 | 16960911 | 2.50E-04 | 3.90E-04 |
| *GATAD2A* | 19 | 19496641 | 19619741 | -6.49 | 8.67E-11 | rs4808203 | 19568659 | 7.20E-12 | 1 | rs3819578 | 19288960 | 0.24 | 0.001 |
| *DHX35* | 20 | 37590980 | 37668366 | 4.68 | 2.93E-06 | rs6065094 | 37453194 | 7.90E-17 | 2.60E-14 | rs6065094 | 37453194 | 7.90E-17 | 2.60E-14 |
| *XPNPEP3* | 22 | 41258260 | 41363888 | 5.36 | 8.21E-08 | rs926914 | 41418154 | 6.90E-10 | 5.50E-04 | rs133047 | 41027819 | 3.50E-09 | 7.60E-06 |
| *NAGA* | 22 | 42454337 | 42466846 | 7.19 | 6.33E-13 | rs134873 | 42657566 | 5.00E-13 | 0.02 | rs5996168 | 42862481 | 1.90E-05 | 1.20E-04 |

Twenty-seven TWAS significant loci, comprising 33 significant TWAS genes, fell outside established (genome-wide significant) schizophrenia risk loci (+/-500 kilobases) as defined by Pardiñas *et al*^5^ (**Table 2**). Of these, 23 loci had a signal attributable to one gene. TWAS significant associations at loci not (yet) implicated in GWAS can arise when multiple variants influence the phenotype through changes in expression, or through the reduced multiple testing burden of gene based analysis^14,16^.

**Table 2.** Twenty-seven TWAS significant loci, containing 33 TWAS significant genes, which fall outside putative schizophrenia risk loci, identified by the Pardiñas et al schizophrenia GWAS, ordered by genomic position. TWAS test statistics for the association between gene expression in the DLPFC and schizophrenia case/control status are denoted by TWAS Z-score and TWAS P-value. Distance (kb) to nearest GWAS loci denotes the genomic distance in kilobases from the gene boundary (Gene Start, Gene End) to the nearest schizophrenia-associated locus, as defined by Pardiñas *et al*. Novel gene associations are highlighted in bold. Novel gene associations where a proximal gene (+/-500 kilobases) has been previously reported are highlighted in bold and with an asterisk. * Note *PCDHA7* and *PCDHAC1* are at the same locus as *NDUFA2* and *PCDHA2*, which have been previously reported in Gusev et al (PMID:29632383), and DNAJA3 is at the same locus as CORO7, which has been previously reported in Gusev et al (PMID:29632383) and Mancuso et al (PMID:28238358).

| Locus | Gene | Chromosome | Gene Start | Gene End | TWAS Z-Score | TWAS P-value | Distance (kb) to nearest GWAS loci |
| --- | --- | --- | --- | --- | --- | --- | --- |
| 1 | **DNAJC11** | 1 | 6694227 | 6761966 | -4.67 | 2.97E-06 | 1630.63 |
| 2 | **LRP8** | 1 | 53708040 | 53793821 | 4.76 | 1.93E-06 | 9570.78 |
| 3 | **PTBP2** | 1 | 97187174 | 97280605 | 4.75 | 2.01E-06 | 519.63 |
| 4 | **KCNN3** | 1 | 154669941 | 154842754 | -5.40 | 6.56E-08 | 4455.77 |
| 5 | *TMEM81* | 1 | 205052256 | 205053588 | 4.55 | 5.40E-06 | 4782.35 |
| 6 | **CGREF1** | 2 | 27322220 | 27341995 | -4.97 | 6.64E-07 | 4568.99 |
| 7 | *LOC100505876* | 2 | 37423634 | 37431886 | 4.87 | 1.09E-06 | 14670.41 |
| 8 | **GABRA2** | 4 | 46251580 | 46392056 | 4.50 | 6.78E-06 | 22808.15 |
| 9 | **SLCO4C1** | 5 | 101569691 | 101632253 | -4.82 | 1.42E-06 | 12821.24 |
| 10 | *NDUFA2* | 5 | 140024947 | 140027370 | -4.81 | 1.53E-06 | 2076.81 |
| 10 | *PCDHA2* | 5 | 140174443 | 140391929 | 5.00 | 5.72E-07 | 2226.30 |
| 10 | **PCDHA7\*** | 5 | 140213968 | 140391929 | 4.64 | 3.55E-06 | 2265.83 |
| 10 | **PCDHAC1\*** | 5 | 140306301 | 140391929 | 4.62 | 3.85E-06 | 2358.16 |
| 11 | **FAM114A2** | 5 | 153371268 | 153418497 | -4.51 | 6.54E-06 | 524.05 |
| 12 | **CUL9** | 6 | 43149921 | 43192325 | 4.54 | 5.50E-06 | 9307.04 |
| 13 | **CALN1** | 7 | 71244475 | 71912136 | 5.12 | 2.98E-07 | 14491.13 |
| 14 | **PXDNL** | 8 | 52232136 | 52722005 | 4.55 | 5.28E-06 | 7753.92 |
| 15 | **MED30** | 8 | 118532964 | 118552501 | -5.23 | 1.74E-07 | 6902.69 |
| 16 | **FAM83H** | 8 | 144806102 | 144815914 | 4.67 | 2.98E-06 | 1447.79 |
| 17 | *TEK* | 9 | 27109146 | 27230172 | -5.16 | 2.44E-07 | 57377.59 |
| 18 | **RASA3** | 13 | 114747193 | 114898095 | 4.44 | 8.89E-06 | 34584.64 |
| 19 | **ZSCAN29** | 15 | 43650369 | 43662258 | -4.85 | 1.22E-06 | 3048.11 |
| 20 | *CORO7* | 16 | 4404542 | 4466962 | -4.84 | 1.27E-06 | 3277.22 |
| 20 | **DNAJA3\*** | 16 | 4475805 | 4506775 | 4.67 | 3.01E-06 | 3237.40 |
| 21 | **COG8** | 16 | 69362523 | 69373526 | 4.59 | 4.45E-06 | 1056.81 |
| 21 | **PDF** | 16 | 69362523 | 69364498 | -4.59 | 4.52E-06 | 1056.81 |
| 22 | **CYB5D2** | 17 | 4046461 | 4060995 | 4.69 | 2.70E-06 | 1830.20 |
| 23 | **ELAC2** | 17 | 12894928 | 12921381 | 4.82 | 1.46E-06 | 4727.79 |
| 24 | **TRIM37** | 17 | 57059999 | 57184266 | 5.51 | 3.67E-08 | 21372.78 |
| 25 | **DCC** | 18 | 49866541 | 51062273 | 4.83 | 1.35E-06 | 1685.42 |
| 26 | *REXO1* | 19 | 1815244 | 1848452 | 4.51 | 6.63E-06 | 10001.28 |
| 27 | **CRELD2** | 22 | 50312282 | 50321186 | 4.49 | 7.19E-06 | 7622.91 |
| 27 | **MAPK11** | 22 | 50702141 | 50708779 | -5.19 | 2.06E-07 | 8012.77 |

The TWAS associations were also significantly correlated with reduced expression at age 18 years, and nominally correlated with increased and reduced expression at two fetal time points (17 and 26 weeks post-conception) in independent samples from BRAINSPAN (**Supplementary Figure 34**).

### Gene-set analysis

We tested ten gene-sets that were reported to be enriched for SNP-association signals in the CLOZUK+PGC2 schizophrenia GWAS^5^. Across these ten sets, two were significantly enriched for TWAS signal; *abnormal long term potentiation* (MP:0002207; p = 6.03×10^-4^) and genes that are intolerant to loss-of-function mutations (*Lek2015 LoFintolerant 90*^21^; p =2.36×10^-3^). However targets of FMRP, one of the most strongly implicated gene-sets in analyses using SNP-level data^5^, showed no evidence for enrichment of TWAS signal (*FMRP_targets*^19^; p = 0.67). This could be in part due to the fact that, in contrast to gene-set analysis of GWAS data which includes data from all measured SNPs within the gene boundaries of each gene-set member, TWAS gene-set analysis only incorporates a subset of SNPs with significant non-zero *cis*-heritable expression in the source tissue, which may reduce the number of informative genes within a given gene-set. Indeed, of the 8 gene-sets enriched for SNP-association signal^5^ which did not demonstrate an enrichment of TWAS signal, 5 were significantly depleted for *cis*-heritable genes in DLPFC (*FMRP targets*^19^; *abnormal behavior*, MP:0004924; *abnormal nervous system electrophysiology*, MP:0002272; *voltage-gated calcium channel complexes*^22^; and *synaptic transmission*, GO:0007268). Targets of FMRP were particularly depleted (p = 3.84×10^-37^), comprising only 3.2% of genes with *cis*-heritable expression in DLPFC compared with 8.9% of genes expressed in that tissue which showed no evidence of *cis*-heritability **(Supplementary Table 5).** However, given the enrichment observed for the much smaller gene-set (i.e. *abnormal long term potentiation*), simple reduction in the number of informative genes cannot provide the sole explanation for loss of signal. Rather, the loss of association for the FMRP targets gene-set suggests at least one of the following alternative hypotheses: a selective loss of signal from schizophrenia associated genes within that dataset; a change in attribution of SNPs to particular genes (in the GWAS analyses, these are based on physical location relative to gene boundaries); being a competitive test, the inclusion of only genes with *cis*-heritable expression as the background set against which specific sets are tested for enrichment might attenuate the signal.

To set the findings in context, and identify novel candidate gene-sets, as in our previous study, we undertook a broader analysis of 2 316 gene-sets which included at least 10 *cis*-heritable genes. These genes represent a TWAS informative subset of a larger data-driven set (N=6 677) described elsewhere^5^. In brief, these sets comprised 101 custom annotated CNS gene-sets^19^; a gene-set of genes intolerant to loss-of-function mutations^21^; 735 gene-sets from Gene Ontology^23,24^ database release 01/02/2016; 663 gene-sets from the fourth ontology level of MGI database version 6^25^; 431 gene-sets from REACTOME^26^ version 55; 203 gene-sets from KEGG^27^ release 04/2015; and 182 gene-sets from OMIM^28^ release 01/02/2016. Significant enrichment after correction for multiple testing (FDR p ≤ 0.05) was found for 5 gene-sets: *nervous system development* (GO:0007399; p = 1.11×10^-4^), *abnormal synaptic transmission*^19^ (p = 0.03), *abnormal long term potentiation*^19^ (p = 0.04), *MGI:reduced long term potentiation* (p = 0.04), and *calcium-dependent cell-cell adhesion* (GO:0016339; p = 0.04). A stepwise multivariable linear modelling approach, where the residual significance of each gene-set was tested after conditioning on the most significant gene-set(s) at each iteration, was used to detect independently significant gene-sets. All but *abnormal long term potentiation* remained significant after conditional analysis (**Supplementary Table 6**).

Genes included in the four conditionally significantly associated gene-sets, and their TWAS association p-values, are provided in **Supplementary Tables 7-10**. Three of the four conditionally independent gene-sets contain a total of 9 TWAS-wide significant genes (**Table 3**, and discussed below); the *calcium-dependent cell-cell adhesion* did not contain any TWAS significant genes, with the signal in this set largely being driven by associations at the proto-cadherin beta gene cluster on chromosome 5 (**Supplementary Table 9**).

**Table 3:** Nine TWAS significant genes which are in gene-sets significantly enriched for schizophrenia TWAS signal. Genes which are not conditionally independent are highlighted using an asterisk (*).

| Gene | Gene-set | Chromosome | Gene Start | Gene Stop | TWAS Z-score | TWAS P-value |
| --- | --- | --- | --- | --- | --- | --- |
| *LRP8* | *abnormal CNS synaptic transmission,*<br>*MGI:reduced long term potentiation* | 1 | 53708040 | 53793821 | 4.76 | 1.93E-06 |
| *CNTN4* | *nervous system development (GO:0007399)* | 3 | 2140549 | 3099645 | 6.03 | 1.65E-09 |
| *GABRA2* | *abnormal CNS synaptic transmission* | 4 | 46251580 | 46392056 | 4.50 | 6.78E-06 |
| *CLCN3* | *abnormal CNS synaptic transmission* | 4 | 170541671 | 170644338 | 5.36 | 8.36E-08 |
| *PCDHA2* | *nervous system development (GO:0007399)* | 5 | 140174443 | 140391929 | 5.00 | 5.72E-07 |
| *PCDHA7\** | *nervous system development (GO:0007399)* | 5 | 140213968 | 140391929 | 4.64 | 3.55E-06 |
| *PCDHAC1* | *nervous system development (GO:0007399)* | 5 | 140306301 | 140391929 | 4.62 | 3.85E-06 |
| *NEURL* | *nervous system development (GO:0007399)* | 10 | 105253734 | 105352309 | -5.69 | 1.31E-08 |
| *CPEB1* | *abnormal CNS synaptic transmission* | 15 | 83211950 | 83316728 | -4.53 | 5.81E-06 |
| *DOC2A* | *abnormal CNS synaptic transmission,*<br>*MGI:reduced long term potentiation,*<br>*nervous system development (GO:0007399)* | 16 | 30016834 | 30024843 | 5.09 | 3.55E-07 |
| *MAPK3* | *abnormal CNS synaptic transmission,*<br>*MGI:reduced long term potentiation* | 16 | 30125425 | 30134630 | 4.56 | 5.19E-06 |

## Discussion

Applying the FUSION method to the largest published schizophrenia GWAS dataset available, we have identified a significant correlation between *cis*-expression in the DLPFC and schizophrenia risk at 89 genes. An earlier FUSION TWAS performed using a smaller schizophrenia GWAS^14^, and the same DLPFC expression weights, identified 44 significant TWAS signals after applying Bonferroni correction, of which 36 remain significant (**Supplementary Table 11**). In the present TWAS, we identified an additional 53 significant associations, 39 of which were neither reported, nor proximal (+/-500kbp) to a significant locus observed in the earlier TWAS based on this tissue. At 32 loci previously associated with schizophrenia at genome-wide levels of significance, we were able to determine that disease risk was correlated to the *cis*-expression of a single gene **(Table 1)** and a further 23 genes outside GWAS loci were similarly implicated as the sole candidates in the region **(Table 2)**. Prioritising putative causal candidate genes and predicted directions of effect at GWAS loci are important steps towards facilitating functional experimentation.

Gene-set analysis of the TWAS data implicated processes related to the development of the CNS, synaptic transmission, and synaptic plasticity through impaired long term potentiation - in particular, processes relevant to glutamatergic and GABAergic transmission (see below). Abnormalities in synaptic transmission and plasticity, and specifically both glutamatergic and GABAergic neurotransmission have been implicated by rare variant studies of schizophrenia^19,20,29–31^, but the findings from common variant studies have, to date, been equivocal. We also identify individual genes within those gene-sets that are TWAS significant; in the context of being individually significantly associated and their membership of biological processes that are implicated in the disorder by the present study, these TWAS significant genes, (discussed below) represent a particularly interesting subset of candidates for generating biological hypotheses.

### Candidates within the abnormal synaptic transmission set and reduced LTP

From the Mouse Genome Informatics (MGI)^32^ set “*abnormal synaptic transmission*” *GABRA2*, *CLCN3*, *DOC2A*, *MAPK3*, *and LRP8* are significantly upregulated (in cases) while *CPEB1* within that set is significantly downregulated in schizophrenia (**Supplementary Table 8**). *DOC2A*, *MAPK3*, *and LRP8* are also members of the MGI *reduced long term potentiation* set **(Supplementary Table 10)**, which was conditionally independently significantly associated in the TWAS gene-set analysis. *GABRA2* encodes the gamma-aminobutyric acid type A (GABA_A_) receptor alpha-2 subunit which combines with other subunits to form pentameric GABA_A_ transmembrane receptors, the main inhibitory receptors in the CNS. A role for altered GABAergic function in schizophrenia has been proposed^33–36^, largely on the basis of imaging studies, animal models of putative intermediate phenotypes, and post-mortem expression studies^37^. In the latter, consistent with our results, elevated immunoreactivity for GABRA2 has been observed in the DLPFC^38^. While this finding has been generally considered to be a post-synaptic homeostatic response to reduced presynaptic GABAergic synthesis and signalling^38^, *in vivo* brain imaging studies studies have not provided support for reduced GABA synthesis^39^. Indeed a recent *in vivo* study of antipsychotic naive patients showing increased GABA in patients challenges this view^40^, as do findings both of no reduction in the major GABA synthetic enzyme GAD67, and an increase in certain types of GABAergic pre-synaptic markers, in post-mortem prefrontal cortex of people with the disorder^41^. Currently, the post-mortem and in vivo imaging data are not decisive in pointing to a primary excess or decrease of GABAergic signalling at glutamatergic neurones in schizophenia^42^. Our finding that elevated *cis*-expression of *GABRA2* is associated with increased risk of schizophrenia suggests the change contributes to pathogenic processes rather than being simply compensatory (see also *CLCN3* below). Although the net effect of over-expression of this receptor on overall GABAergic function is unclear, our findings suggests the possibility that targeting treatments at antagonism at this receptor complex might be more appropriate than agonism, as has largely been the focus to date. At a wider level, this finding adds further genetic support for the GABAergic hypothesis of schizophrenia and complements a recent study which reported enrichment for components of GABA_A_ receptor complexes in CNVs in people with schizophrenia^19^.

*CLCN3* encodes an intracellular, voltage-gated chloride channel. It is involved in regulating neurotransmitter vesicle turnover at both excitatory glutamatergic and inhibitory GABAergic synapses, can be pre- or postsynaptic, and is thought to be an important regulator of synaptic plasticity^43^ ^44^. Neurotransmitter release at inhibitory GABAergic synapses is reduced in CLCN3-null mice while glutamate release is increased ^43^ ^44^; our finding, and that of another study based on the SMR approach^17^, of increased expression of this gene might suggest the reverse effect (reduced glutamate, increased GABA) in schizophrenia consistent with the aberrance of both excitatory and inhibitory signalling in schizophrenia.

*DOC2A* and *MAPK3*, encoding double C2 domain alpha and mitogen-activated protein kinase 3 respectively, map to 16p11.2, duplication of which is associated with schizophrenia^45^. Both are *cis*-upregulated in schizophrenia in the present study and in a previous TWAS study^14^, consistent with the possibility that one or both contribute to the pathogenicity of the 16p11.2 duplication. Evidence from animal studies supports the hypothesis that both *MAPK3*^14^ and *DOC2A*^46^ can contribute to CNS phenotypes; in the present study, conditional analysis suggests expression of both genes contributes to the association signal at this locus, the conditional evidence being much stronger for expression of *DOC2A* (conditional on *MAPK3*, p=3.5×10^-7^) than *MAPK3* (conditional on *DOC2A* p=0.014). Reduction in *DOC2A* expression results in a reduction of spontaneous glutamate release from hippocampal neurons in culture, and increases glutamatergic synaptic efficiency; and synaptic efficacy^47^, suggesting the possibility that the increased *cis*-expression in schizophrenia predicted by the TWAS might result in hypofunction at glutamatergic synapses. Evidence implicating *MAPK3* in schizophrenia is discussed in an earlier TWAS study^14^.

*CPEB1*, (reduced *cis*-expression) encoding cytoplasmic polyadenylation element binding protein 1, is expressed in dendrites where it facilitates activity dependent mRNA translation^48,49^ and memory consolidation^50^. It has been reported to bind, and act antagonistically to, FMRP which is a translational repressor^50^. Given that people with schizophrenia are enriched for loss-of-function mutations in genes encoding targets of FMRP^20,30^, we postulate the relative reduction of expression of CPEB1, a facilitator of translation of those genes, might similarly increase risk of schizophrenia by reduced expression of FMRP targets.

The other gene in the set is *LRP8* (*cis*-upregulation) which encodes LDL Receptor Related Protein 8. *LRP8* is widely expressed, but is particularly highly expressed in the developing foetal brain^51^. A previous candidate gene study has reported (non-genomewide significant) association between *LRP8* and a combined schizophrenia-bipolar phenotype^51^. Moreover, as in this study, risk alleles were associated (non-genomewide significant) with higher *LRP8* mRNA expression in brain^51^. *LRP8* is a component of the NMDA receptor complex (NMDA-R) at glutamatergic synapses^52^, a complex repeatedly implicated in schizophrenia^19,29,30^. It is also the major receptor, and effector, of the actions of reelin as a modulator of plasticity, memory and learning in the adult brain^53^. It should be noted that *LRP8* has non-synaptic functions of credible relevance to schizophrenia, including mediating reelin related functions on neuronal migration, neurogenesis, and neuronal differentiation^54^. However, these processes are not implicated by the gene-set analysis. Moreover, it is unclear how increased expression of *LRP8* might relate to the reduced reelin function that has most commonly been linked to schizophrenia.

Although a member of the gene-set *nervous system development (GO:0007399)*, *NEURL1* (*cis*-downregulated), which encodes neuralized E3 ubiquitin protein ligase 1, is functionally better characterized as gene involved in reduction of LTP and synaptic plasticity. *NEURL1* is widely expressed in brain. Its mRNA is present in dendrites where translation is synaptic activity dependent^55^. Suppression of *NEURL1* reduces glutamatergic AMPA receptor expression and impairs both LTP and LTD forms of synaptic plasticity and memory formation. Overexpression has the opposite effects^55^. Synaptic activity induced *NEURL1* expression is thought to increase the number of AMPA receptors, and the number of functional AMPA containing synapses, via activating CPEB3, a protein which like another TWAS significant member of the same family CBEP1 (above) is a facilitator of local mRNA translation^55^.

### Candidates within the nervous system development (GO:0007399) gene-set

TWAS significant genes within this gene-set include *CNTN4*, *PCDHA2*, *PCDHA7*, *PCDHAC1* **(Supplementary Table 7)** as well as *NEURL1* and *DOC2A* which were discussed in the previous section. *CNTN4* (*cis*-upregulated) encodes contactin 4, a member of the contactin family of axon-associated cell adhesion molecules. Similar findings with *CNTN4* were previously reported^17^ in a study which went on to show that overexpression of this gene in zebrafish resulted in smaller head size and a reduction in cellular proliferation. *CNTN4* is most highly expressed in the CNS, particularly in pyramidal neurons and interneurons; while it is proposed to pay a role in neurite growth and axonal targeting, its range of actions and the mechanisms underpinning them are poorly understood^56^. The associated protocadherins (*PCDHA2*, *PCDHA7*, *PCDHAC1*, all upregulated) map to one of several protocadherin clusters on chromosome 5. They are physically overlapping, and therefore although *PCDHA2*, and *PCDHAC1* are conditionally independent (**Supplementary Table 3**), we consider all three genes to represent a single association signal that affects multiple functionally related genes, likely including members of the beta protocadherin family as well (**Supplementary Table 9**). *PCDHA* genes, which were originally referred to as cadherin related neuronal receptors, are largely localized in dendritic/synaptic structures^57^. They have been poorly functionally characterized, but are involved at many stages of neural circuitry development including axon outgrowth, and the formation of complex dendritic structures and postsynaptic spines^58^. Most studies have observed impaired synaptic development and function after knockout or suppression of PCDHA genes, for example knock out of *PCDHA2*, whose expression is largely confined to serotonergic and noradrenergic neurons, selectively impairs serotonergic neuronal projections and function, and induces multiple depression related phenotypes in mice^59^. The impact of overexpression of *PCDHA* genes has not been reported. However, one study^60^ reported that overexpression of some types of protocadherins (gamma) impairs synaptic development by disrupting the activity of *neurexin 1* (deletion CNVs of which are associated with schizophrenia^29^), so it seems therefore seems that over- or under-expression of individual members of this complex superfamily of genes impacts on CNS development.

The present study represents one approach to mine GWAS association findings, but it is not without limitations. Here, as others have done before in smaller datasets and/or with different methods, we show that changes in gene expression can be imputed into case-control studies, but such data are essentially correlational and do not prove those changes are causal. Nevertheless, as noted above, even when prior hypotheses are not imposed on the dataset, (i.e. data driven analyses), our findings show evidence for convergence with earlier rare variant studies, albeit such convergence is largely at the level of systems and processes rather than specific genes. Whether this reflects a true lack of overlap between common variant and rare variant studies is at present unclear given the low power of rare variant studies to date, and the modest power both of the eQTL and GWAS datasets. In this respect, the relative depletion of some of the major common and rare variant associated gene-sets (LOF intolerant and targets of FMRP) for *cis*-heritable expression is notable. We postulate this may suggest that, just as for damaging mutations, selection pressures constrain the impact of *cis*-eQTLs to quite small effects that will require much larger eQTL datasets to detect. Beyond sample size, the tissue source for the eQTLs may also be an important limitation. The present study is based upon *post-mortem* adult human brain tissue from the DLPFC, as we judge this to be optimal in terms of the balance of sample size and pathogenic plausibility. However, under the hypothesis that at least some of the processes that underpin schizophrenia exert their pathogenic effects during early development, it will be important to evaluate eQTL datasets from foetal samples as these become available.

In summary, we have undertaken a TWAS study exploiting the largest published schizophrenia GWAS dataset to date. We identify large numbers of novel genes whose expression correlate with schizophrenia, implicate processes of high biological plausibility related to synaptic function, and identify candidate genes to which we assign predicted directions of effect which should facilitate experimental studies geared towards a better mechanistic understanding of schizophrenia pathogenesis.

## Acknowledgements

This work was funded by Medical Research Council (MRC) Centre (MR/L010305/1), Program Grant (G0800509) and Project Grants (MR/L011794/1). Additional support was provided by National Institute of Mental Health (NIMH) Psychiatric Genomics Consortium (PGC) Grant (5U01MH109514-02). We acknowledge Cardiff University MRC Centre HPC team (Mark Einon, David Simpson, Ellis Pires), and Cardiff University Advanced Research Computing division (Wayne Lawrence), for support with the use and setup of computational infrastructures.

## References

1. Addington, J. & Heinssen, R. Prediction and prevention of psychosis in youth at clinical high risk. Annu. Rev. Clin. Psychol. 8, 269–289 (2012).

2. Nordentoft, M., Mortensen, P. B. & Pedersen, C. B. Absolute risk of suicide after first hospital contact in mental disorder. Arch. Gen. Psychiatry 68, 1058–1064 (2011).

3. Owen, M. J., Sawa, A. & Mortensen, P. B. Schizophrenia. Lancet 388, 86–97 (2016).

4. Chesney, E., Goodwin, G. M. & Fazel, S. Risks of all-cause and suicide mortality in mental disorders: a meta-review. World Psychiatry 13, 153–160 (2014).

5. Pardiñas, A. F. et al. Common schizophrenia alleles are enriched in mutation-intolerant genes and in regions under strong background selection. Nat. Genet. (2018). doi:10.1038/s41588-018-0059-2

6. Singh, T. et al. Rare loss-of-function variants in SETD1A are associated with schizophrenia and developmental disorders. Nat. Neurosci. 19, 571–577 (2016).

7. Rees, E. et al. Analysis of copy number variations at 15 schizophrenia-associated loci. Br. J. Psychiatry 204, 108–114 (2014).

8. Pasaniuc, B. & Price, A. L. Dissecting the genetics of complex traits using summary association statistics. Nat. Rev. Genet. 18, 117–127 (2017).

9. Richards, A. L. et al. Schizophrenia susceptibility alleles are enriched for alleles that affect gene expression in adult human brain. Mol. Psychiatry 17, 193–201 (2012).

10. Schizophrenia Working Group of the Psychiatric Genomics Consortium. Biological insights from 108 schizophrenia-associated genetic loci. Nature 511, 421–427 (2014).

11. Zhu, Z. et al. Integration of summary data from GWAS and eQTL studies predicts complex trait gene targets. Nat. Genet. 48, 481–487 (2016).

12. Gamazon, E. R. et al. A gene-based association method for mapping traits using reference transcriptome data. Nat. Genet. 47, 1091–1098 (2015).

13. Gusev, A. et al. Integrative approaches for large-scale transcriptome-wide association studies. Nat. Genet. 48, 245–252 (2016).

14. Gusev, A. et al. Transcriptome-wide association study of schizophrenia and chromatin activity yields mechanistic disease insights. Nat Genet 067355 (2018).

15. Barbeira, A. N. et al. Exploring the phenotypic consequences of tissue specific gene expression variation inferred from GWAS summary statistics. Nat. Commun. 9, 1825 (2018).

16. Mancuso, N. et al. Integrating Gene Expression with Summary Association Statistics to Identify Genes Associated with 30 Complex Traits. Am. J. Hum. Genet. 100, 473–487 (2017).

17. Fromer, M. et al. Gene expression elucidates functional impact of polygenic risk for schizophrenia. Nat. Neurosci. 19, 1442–1453 (2016).

18. R Core Team. R: A Language and Environment for Statistical Computing. (2017).

19. Pocklington, A. J. et al. Novel Findings from CNVs Implicate Inhibitory and Excitatory Signaling Complexes in Schizophrenia. Neuron 86, 1203–1214 (2015).

20. Genovese, G. et al. Increased burden of ultra-rare protein-altering variants among 4,877 individuals with schizophrenia. Nat. Neurosci. 19, 1433–1441 (2016).

21. Lek, M. et al. Analysis of protein-coding genetic variation in 60,706 humans. Nature 536, 285–291 (2016).

22. Müller, C. S. et al. Quantitative proteomics of the Cav2 channel nano-environments in the mammalian brain. Proc. Natl. Acad. Sci. U. S. A. 107, 14950–14957 (2010).

23. Network and Pathway Analysis Subgroup of Psychiatric Genomics Consortium. Psychiatric genome-wide association study analyses implicate neuronal, immune and histone pathways. Nat. Neurosci. 18, 199–209 (2015).

24. Gene Ontology Consortium. Gene Ontology Consortium: going forward. Nucleic Acids Res. 43, D1049–56 (2015).

25. Blake, J. A. et al. The Mouse Genome Database: integration of and access to knowledge about the laboratory mouse. Nucleic Acids Res. 42, D810–7 (2014).

26. Fabregat, A. et al. The Reactome pathway Knowledgebase. Nucleic Acids Res. 44, D481–7 (2016).

27. Kanehisa, M., Sato, Y., Kawashima, M., Furumichi, M. & Tanabe, M. KEGG as a reference resource for gene and protein annotation. Nucleic Acids Res. 44, D457–62 (2016).

28. Amberger, J. S., Bocchini, C. A., Schiettecatte, F., Scott, A. F. & Hamosh, A. OMIM.org: Online Mendelian Inheritance in Man (OMIM®), an online catalog of human genes and genetic disorders. Nucleic Acids Res. 43, D789–98 (2015).

29. Kirov, G. et al. De novo CNV analysis implicates specific abnormalities of postsynaptic signalling complexes in the pathogenesis of schizophrenia. Mol. Psychiatry 17, 142–153 (2012).

30. Fromer, M. et al. De novo mutations in schizophrenia implicate synaptic networks. Nature 506, 179–184 (2014).

31. CNV and Schizophrenia Working Groups of the Psychiatric Genomics Consortium & Psychosis Endophenotypes International Consortium. Contribution of copy number variants to schizophrenia from a genome-wide study of 41,321 subjects. Nat. Genet. 49, 27–35 (2017).

32. Smith, C. L. et al. Mouse Genome Database (MGD)-2018: knowledgebase for the laboratory mouse. Nucleic Acids Res. 46, D836–D842 (2018).

33. Olney, J. W. & Farber, N. B. Glutamate receptor dysfunction and schizophrenia. Arch. Gen. Psychiatry 52, 998–1007 (1995).

34. Lewis, S. Synaptic transmission: A closer look at presynaptic GABA(B) receptors. Nat. Rev. Neurosci. 11, 664 (2010).

35. Roberts, E. Prospects for research on schizophrenia. An hypotheses suggesting that there is a defect in the GABA system in schizophrenia. Neurosci. Res. Program Bull. 10, 468–482 (1972).

36. Lewis, D. A., Hashimoto, T. & Volk, D. W. Cortical inhibitory neurons and schizophrenia. Nat. Rev. Neurosci. 6, 312–324 (2005).

37. Inan, M., Petros, T. J. & Anderson, S. A. Losing your inhibition: linking cortical GABAergic interneurons to schizophrenia. Neurobiol. Dis. 53, 36–48 (2013).

38. Stan, A. D. & Lewis, D. A. Altered cortical GABA neurotransmission in schizophrenia: insights into novel therapeutic strategies. Curr. Pharm. Biotechnol. 13, 1557–1562 (2012).

39. Egerton, A., Modinos, G., Ferrera, D. & McGuire, P. Neuroimaging studies of GABA in schizophrenia: a systematic review with meta-analysis. Transl. Psychiatry 7, e1147 (2017).

40. de la Fuente-Sandoval, C. et al. Prefrontal and Striatal Gamma-Aminobutyric Acid Levels and the Effect of Antipsychotic Treatment in First-Episode Psychosis Patients. Biol. Psychiatry 83, 475–483 (2018).

41. Rocco, B. R., DeDionisio, A. M., Lewis, D. A. & Fish, K. N. Alterations in a Unique Class of Cortical Chandelier Cell Axon Cartridges in Schizophrenia. Biol. Psychiatry 82, 40–48 (2017).

42. de Jonge, J. C., Vinkers, C. H., Hulshoff Pol, H. E. & Marsman, A. GABAergic Mechanisms in Schizophrenia: Linking Postmortem and In Vivo Studies. Front. Psychiatry 8, 118 (2017).

43. Guzman, R. E., Alekov, A. K., Filippov, M., Hegermann, J. & Fahlke, C. Involvement of ClC-3 chloride/proton exchangers in controlling glutamatergic synaptic strength in cultured hippocampal neurons. Front. Cell. Neurosci. 8, 143 (2014).

44. Riazanski, V. et al. Presynaptic CLC-3 determines quantal size of inhibitory transmission in the hippocampus. Nat. Neurosci. 14, 487–494 (2011).

45. McCarthy, S. E. et al. Microduplications of 16p11.2 are associated with schizophrenia. Nat. Genet. 41, 1223–1227 (2009).

46. McCammon, J. M., Blaker-Lee, A., Chen, X. & Sive, H. The 16p11.2 homologs fam57ba and doc2a generate certain brain and body phenotypes. Hum. Mol. Genet. 26, 3699–3712 (2017).

47. Ramirez, D. M. O. et al. Loss of Doc2-Dependent Spontaneous Neurotransmission Augments Glutamatergic Synaptic Strength. J. Neurosci. 37, 6224–6230 (2017).

48. Hake, L. E. & Richter, J. D. CPEB is a specificity factor that mediates cytoplasmic polyadenylation during Xenopus oocyte maturation. Cell 79, 617–627 (1994).

49. Yue, J. et al. Anxiolytic effect of CPEB1 knockdown on the amygdala of a mouse model of inflammatory pain. Brain Res. Bull. 137, 156–165 (2018).

50. Udagawa, T. et al. Genetic and acute CPEB1 depletion ameliorate fragile X pathophysiology. Nat. Med. 19, 1473–1477 (2013).

51. Li, M. et al. Convergent Lines of Evidence Support LRP8 as a Susceptibility Gene for Psychosis. Mol. Neurobiol. 53, 6608–6619 (2016).

52. Hoe, H.-S. et al. Apolipoprotein E receptor 2 interactions with the N-methyl-D-aspartate receptor. J. Biol. Chem. 281, 3425–3431 (2006).

53. Telese, F. et al. LRP8-Reelin-Regulated Neuronal Enhancer Signature Underlying Learning and Memory Formation. Neuron 86, 696–710 (2015).

54. Santana, J. & Marzolo, M.-P. The functions of Reelin in membrane trafficking and cytoskeletal dynamics: implications for neuronal migration, polarization and differentiation. Biochem. J 474, 3137–3165 (2017).

55. Pavlopoulos, E. et al. Neuralized1 activates CPEB3: a function for nonproteolytic ubiquitin in synaptic plasticity and memory storage. Cell 147, 1369–1383 (2011).

56. Oguro-Ando, A., Zuko, A., Kleijer, K. T. E. & Burbach, J. P. H. A current view on contactin-4, -5, and -6: Implications in neurodevelopmental disorders. Mol. Cell. Neurosci. 81, 72–83 (2017).

57. Keeler, A. B., Molumby, M. J. & Weiner, J. A. Protocadherins branch out: Multiple roles in dendrite development. Cell Adh. Migr. 9, 214–226 (2015).

58. Peek, S. L., Mah, K. M. & Weiner, J. A. Regulation of neural circuit formation by protocadherins. Cell. Mol. Life Sci. 74, 4133–4157 (2017).

59. Chen, W. V. et al. Pcdhαc2 is required for axonal tiling and assembly of serotonergic circuitries in mice. Science 356, 406–411 (2017).

60. Molumby, M. J. et al. γ-Protocadherins Interact with Neuroligin-1 and Negatively Regulate Dendritic Spine Morphogenesis. Cell Rep. 18, 2702–2714 (2017).

